# Disparate Bone Anabolic Cues Activate Bone Formation by Regulating the Rapid Lysosomal Degradation of Sclerostin Protein

**DOI:** 10.1101/2020.10.26.355800

**Authors:** Nicole R. Gould, Katrina M. Williams, Humberto C. Joca, Olivia M. Torre, James S. Lyons, Jenna M. Leser, Manasa P. Srikanth, Marcus Hughes, Ramzi J. Khairallah, Ricardo A. Feldman, Christopher W. Ward, Joseph P. Stains

**Affiliations:** Department of Orthopaedics, University of Maryland School of Medicine, Baltimore, MD, 21201, USA; Center for Biomedical Engineering and Technology, University of Maryland School of Medicine, Baltimore, MD, 21201, USA; Department of Microbiology and Immunology, University of Maryland School of Medicine, Baltimore, MD, 21201, USA; Myologica, LLC, New Market, MD 21212, USA

## Abstract

The down regulation of sclerostin mediates bone formation in response to mechanical cues and parathyroid hormone (PTH). To date, the regulation of sclerostin has been attributed exclusively to the transcriptional downregulation that occurs hours after stimulation. Here, we describe, for the first time, the rapid post-translational degradation of sclerostin protein by the lysosome following mechanical load or PTH. We present a unifying model, integrating both new and established mechanically- and hormonally-activated effectors into the regulated degradation of sclerostin by lysosomes. Using an *in vivo* mechanical loading model, we find transient inhibition of lysosomal degradation or the upstream mechano-signaling pathway controlling sclerostin abundance impairs subsequent load-induced bone formation. We also link dysfunctional lysosomes to aberrant sclerostin regulation using Gaucher disease iPSCs. These results inform a paradigm shift in how bone anabolic cues post-translationally regulate sclerostin and expands our understanding of how osteocytes regulate this fundamentally important protein to regulate bone formation.

## Introduction

The osteocyte derived protein, sclerostin, is a fundamentally important inhibitor of bone formation, with decreases in abundance mediating mechanically- and hormonally-induced bone formation. Sclerostin (gene name *Sost*) is a secreted 27 kDa glycoprotein that inhibits the differentiation and activity of bone-forming osteoblasts by antagonizing the Wnt signaling pathway (1, 2). Genetic deletion of the *Sost* gene in mice results in extraordinarily high bone mass (3). In humans, mutations in the *SOST* gene underlie high bone mass and bone overgrowth in patients with sclerosteosis and van Buchem disease (4-6). Accordingly, regulating sclerostin bioavailability has tremendous therapeutic potential for conditions of low bone mass, such as osteoporosis. Indeed, targeting sclerostin protein with neutralizing antibodies is incredibly effective at increasing bone mass, and Romosozumab, a humanized monoclonal antibody targeting sclerostin, has been FDA approved to treat osteoporosis in post-menopausal women at a high risk for fracture (7, 8). Despite this critical role for sclerostin in skeletal homeostasis and its therapeutic potential, there are substantial gaps with respect to the molecular control of this key regulatory protein.

In response to bone mechanical loading, osteocytes sense and respond to fluid shear stress (FSS) in the lacunar canalicular network by ultimately decreasing sclerostin protein abundance, unleashing osteoblast differentiation and bone formation. Likewise, parathyroid hormone (PTH) also works through decreasing sclerostin abundance (9, 10). When administered intermittently, PTH causes net bone formation, which has been exploited in the clinic through the established osteoanabolic drug, teriparatide (PTH, amino acids 1-34). Despite their clinical application, little is known about how mechanical load and PTH exposure, two disparate bone anabolic signals, regulate sclerostin protein bioavailability. To date, the regulation of sclerostin abundance has been exclusively attributed to the transcriptional downregulation of the *Sost* gene that occurs hours after mechanical load or PTH exposure (11, 12).

Using a recently established osteocyte-like cell line Ocy454 cells, which is one of the few cell lines that reliably express detectable sclerostin protein (13, 14), we described a mechano-transduction pathway that regulates osteocyte sclerostin protein abundance in response to FSS *in vitro* (Fig. 1A) (15, 16). Using this *in vitro* model, we previously found that osteocyte mechano-signaling required a subset of detyrosinated microtubules, which transduce load signals to activate NADPH Oxidase 2 (NOX2), which produces reactive oxygen species (ROS) signals that elicit a Transient Receptor Potential Vanilloid 4 (TRPV4)-dependent primary calcium (Ca^2+^) influx. Calcium/calmodulin-dependent kinase II (CaMKII) is activated in response to this primary Ca^2+^ influx and is required for reduction of osteocyte sclerostin protein abundance (Fig. 1A). While these discoveries integrated with and extended several established models of the osteocyte mechanical response (17-20), we found the loss of sclerostin protein was surprisingly rapid and was likely wholly distinct from the well-characterized transcriptional regulation of the *Sost* gene, which occurs on the hours timescale (11, 12). Despite its physiologic significance, surprisingly little is known about the post-translational control of sclerostin protein. Additionally, given the *in vitro* nature of our prior work on this pathway, the contribution of this mechano-transduction pathway to *in vivo* bone mechano-responsiveness remained unresolved. Here, we examined the mechanism regulating the rapid decline in osteocyte sclerostin protein, its relevance *in vivo* in bone physiology and skeletal disease, and extended this rapid loss of sclerostin to PTH, another clinically relevant bone anabolic signal.

**Fig. 1:**
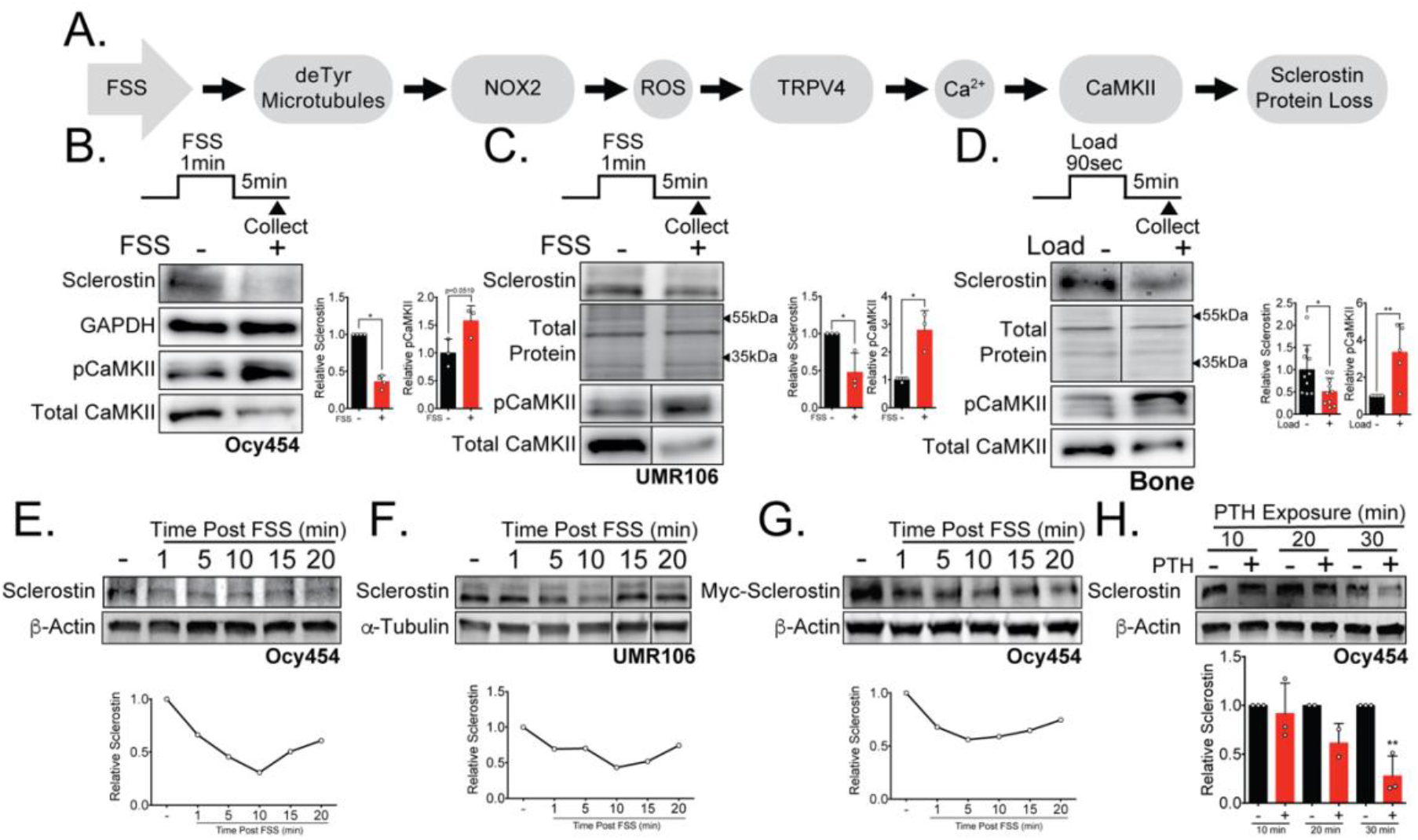
Sclerostin protein is rapidly degraded after anabolic stimuli in vitro and in vivo. **(A)** FSS causes the rapid loss of sclerostin protein through a number of molecular mediators. **(B)** Ocy454 cells (n=3-4) or **(C)** UMR106 cells (n=3) were exposed to one minute of FSS at 4 dynes/cm^2^ and lysed five minutes post-flow. Western blots were probed for sclerostin, GAPDH, pCAMKII, and total CaMKII. For each antibody, blots are from a single gel and exposure; a vertical black line indicates removal of irrelevant lanes. **(D)** 16-week old female C57Bl/6 mice were ulnar loaded (2.8N, 90sec, 2Hz), cortical osteocyte enriched lysates isolated 5 minutes post-load, and western blots probed for sclerostin, pCaMKII, and total CaMKII (n=5 independent animals). Sclerostin abundance relative to the loading control or pCaMKII relative to total CaMKII was quantified. For each antibody, blots are from a single gel and exposure; a vertical black line indicates removal of irrelevant lanes. **(E)** Ocy454 cells with endogenous sclerostin (n=2), **(F)** UMR106 cells with endogenous sclerostin (n=4) or, **(G)** Ocy454 cells transfected with Myc-tagged sclerostin (n=1) were subjected to five minutes of FSS at 4 dynes/cm^2^ and lysed at the indicated times post-flow. Western blots were probed for sclerostin and β-actin. A representative time course is shown for each. Sclerostin abundance relative to the loading control was quantified. For each antibody, blots are from a single gel and exposure; a vertical black line indicates removal of irrelevant lanes. **(H)** Ocy454 cells transfected with GFP-sclerostin were treated with vehicle or PTH (1-34) (10nM) for the indicated time and were lysed at time indicated. Western blots were probed for sclerostin and β-actin (n=2-3). Graphs depict mean ± SD. *p<0.05, **p<0.01 by two-tailed t-tests (A-C) or two-way ANOVA with Holm-Sidak post-hoc correction (H).

## Results

### Sclerostin protein is rapidly lost after mechanical load *in vitro* and *in vivo*

To characterize the dynamics of mechanically stimulated sclerostin protein loss in osteocytes, we examined two sclerostin expressing cell lines, Ocy454 osteocytes and UMR106 osteosarcoma cells. Consistent with our previous work (15, 16), we observed the mechano-activated increase in CaMKII phosphorylation and decrease in sclerostin protein abundance in both Ocy454 osteocytes (Fig. 1B) and UMR106 osteosarcoma cells (Fig. 1C) within 5 minutes of the acute application of FSS. Next, we examined if *in vivo* loading mimicked the *in vitro* kinetics of rapid sclerostin loss. Sclerostin protein abundance was assessed in 16-week-old female mice subjected to a single bout of ulnar load. Ulnae were harvested 5 minutes post-load and osteocyte-enriched cortical bone lysates were profiled by western blotting. As occurred *in vitro*, sclerostin protein abundance was reduced and CaMKII activated in loaded versus contralateral non-loaded limbs (Fig. 1D), establishing that rapid sclerostin downregulation also occurs *in vivo* following a single bout of mechanical stimulus.

A temporal assessment of sclerostin protein abundance following FSS in Ocy454 cells (Fig. 1E) and UMR106 cells (Fig. 1F) revealed a decrease in sclerostin protein as early as 1-minute post FSS, with the nadir of protein expression occurring at 10 minutes, followed by a subsequent rebound. This rapid regulation strongly suggested post-translational control of sclerostin abundance. Consistent with the possibility of post-translational control of sclerostin, we recently reported that a single 5-minute bout of FSS, as used here, does not decrease *Sost* mRNA levels despite a decrease in sclerostin protein (16). To confirm the post-translational control of sclerostin, Ocy454 cells were transfected with either a myc- or GFP-tagged sclerostin expression vector under control of a CMV promoter, which is not regulated by the same transcriptional elements that regulate *Sost* transcription. Similar to endogenous sclerostin, overexpressed sclerostin protein was rapidly and transiently decreased by FSS (Fig. 1G and Supplemental Fig. 1A). The regulated degradation did not extend to all proteins as pro-collagen type 1α1 abundance was unchanged following FSS in Ocy454 cells (Supplemental Fig. 1B), supporting at least some level of specificity for the regulated degradation of sclerostin.

### PTH1-34 treatment also causes rapid sclerostin protein loss

Like mechanical loading, intermittent administration of PTH stimulates bone remodeling and results in net bone formation (21). Both chronic and intermittent PTH treatment decrease osteocyte *Sost* gene expression hours after exposure (11); however, rapid loss of sclerostin protein has not been reported. In Ocy454 cells, sclerostin protein abundance decreased after 30 minutes of PTH treatment (Supplemental Fig. 1C) with no change in pro-collagen type 1α1 (Supplemental Fig. 1D). Examining the post-translational regulation of sclerostin protein abundance in GFP-sclerostin transfected Ocy454 cells, we show PTH treatment caused sclerostin loss over a 30-minute time period (Fig. 1H). While the kinetics of PTH were slower than FSS, the loss of sclerostin protein was still relatively rapid and occurred with overexpressed protein, supporting that both mechanical load and PTH regulates post-translational control of sclerostin in close temporal relation to application of the bone anabolic cue.

### Sclerostin is degraded through the lysosome after exposure to bone anabolic stimuli

The post-translational loss of sclerostin protein shortly after exposure to a bone anabolic stimulus likely occurs through either rapid degradation or secretion. Accordingly, we interrogated the route of rapid sclerostin loss after anabolic stimuli by blocking degradation or secretion pathways using different pharmacological inhibitors. In these degradation assays, cells were treated with cycloheximide to prevent *de novo* protein synthesis, as well as either the lysosome inhibitor, Bafilomycin A1, the proteasome inhibitor, MG-132, or the secretion inhibitor, Brefeldin A. Subsequently, the cells were exposed to 5 minutes of FSS or 30 minutes of PTH, and sclerostin protein abundance monitored by western blot. In Ocy454 cells transfected with GFP-sclerostin, inhibition of lysosomal function with Bafilomycin A1 prevented both FSS- and PTH-induced degradation of sclerostin protein (Fig. 2A and 2B), whereas inhibition of proteasomal degradation with MG-132 or secretion with Brefeldin A had no effect on the FSS-induced decrease in sclerostin (Fig. 2A). Likewise, inhibition of lysosomal activity with Bafilomycin A1 or leupeptin prevented FSS-induced sclerostin degradation in UMR106 osteosarcoma cells (Supplemental Fig. 2A). Notably, in the absence of mechanical stimulus, the half-life of sclerostin protein is about 3 hours, in contrast to the minutes scale observed after stimulation, confirming that the kinetics of sclerostin degradation are altered in response to bone anabolic cues (Supplemental Fig. 2B).

**Fig. 2:**
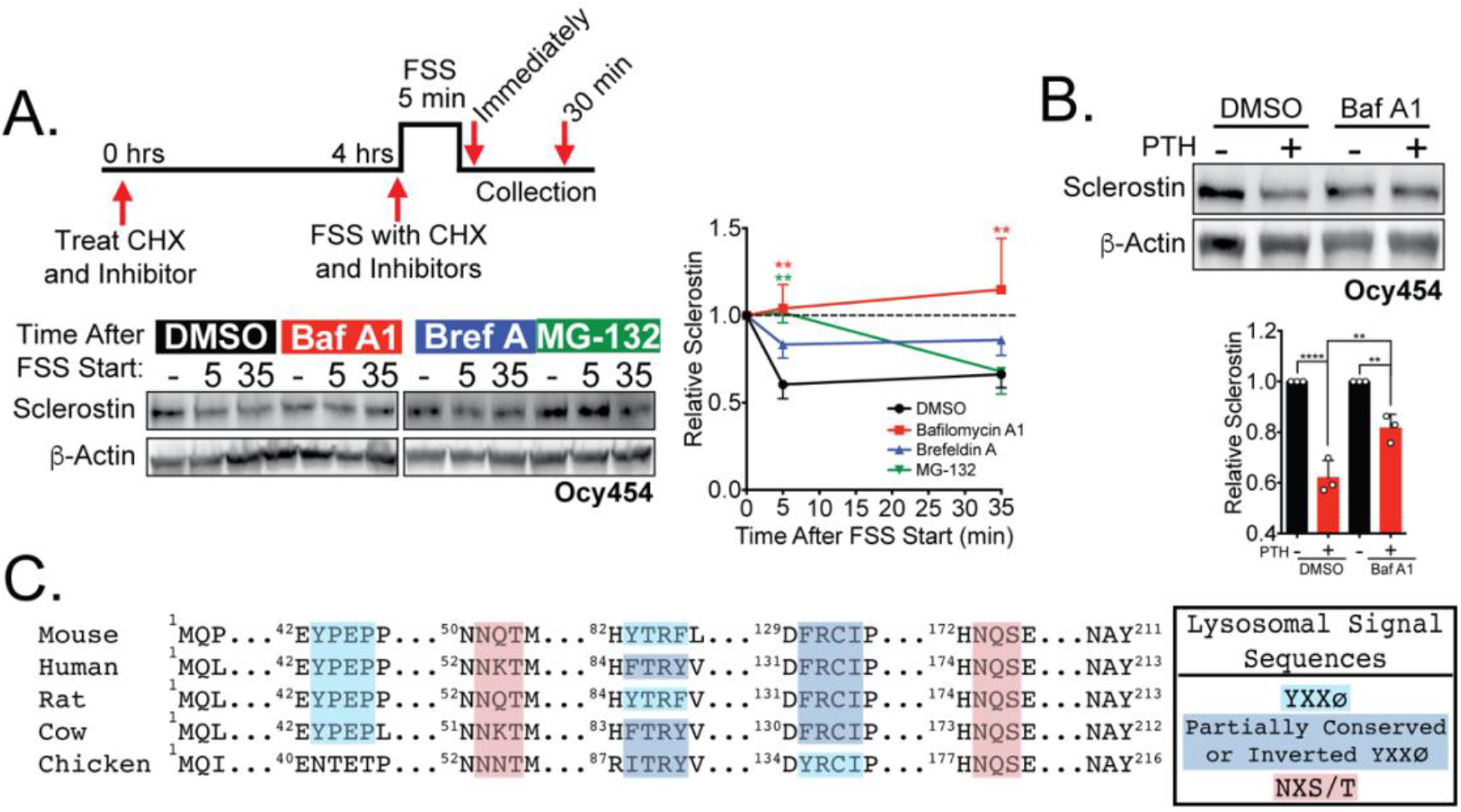
Sclerostin is rapidly degraded by the lysosome following bone anabolic stimuli. **(A)** Ocy454 cells transfected with GFP-sclerostin were treated with cycloheximide (150µg/mL) and either DMSO (0.1%), Bafilomycin A1 (100nM), Brefeldin A (2μm), or MG-132 (10μm) 4 hours prior to FSS. Cells were subjected to 5 minutes of FSS at 4 dynes/cm^2^ and lysed immediately after the end of FSS or 30 minutes after the conclusion of FSS. Western blots were probed for sclerostin and β-actin. Timecourse shows mean ± SEM (n=3-6 independent experiments/group). B)Ocy454 cells transfected with GFP-sclerostin were pre-treated with DMSO (0.1%) or Bafilomycin A1 (100nM) for 30 minutes prior to the addition of vehicle or PTH (1-34) (10nM) for an additional 30 minutes (n=3). Sclerostin abundance relative to the loading control was quantified. Graph depicts mean ± SD. *p<0.05, **p<0.01, ****p<0.0001 by two-way ANOVA with Holm-Sidak post-hoc correction. **(C)** Amino acid sequences for sclerostin from mouse, human, rat, cow, and chicken were aligned using NCBI COBALT. Abbreviated sequences are shown and are annotated for putative lysosomal signal sequences. Full sequences are presented in Supplemental Fig. 2C.

In support of this notion of lysosomal degradation of sclerostin, many secreted N-linked mannose-6-phosphate and GlcNAC modified glycoproteins, like sclerostin (22), are targeted to the lysosome to regulate abundance (23). Typically, these proteins contain conserved lysosomal signal sequences, particularly the Asn-X-Ser/Thr sites that are subjected to N-linked glycosylation (23). *In silico* analysis of the sclerostin protein amino acid sequence identified at least two putative sites for N-linked glycosylation (Asn-X-Ser/Thr) and three Tyr-X-X-Φ lysosome targeting motifs that are conserved across species (Fig. 2C, Supplemental Fig. 2C).

### Sclerostin protein co-localizes with lysosomal markers

To validate that sclerostin is targeted to the lysosome, we examined the sub-cellular distribution of sclerostin and lysosomes in cultured Ocy454 cells. Both endogenous and exogenous sclerostin were found in distinct puncta in Ocy454 cells (Fig. 3A). In live Ocy454 cells, GFP-sclerostin was co-localized with acidic vesicles identified by LysoTracker and significantly co-localized with lysosomes labeled by siR-lysosome (Fig. 3B). In fixed cells, endogenous sclerostin co-localized with p62/sequestosome-1 protein, an autophagy cargo adapter protein that shuttles proteins for lysosomal degradation (Fig. 3C). That sclerostin is found localized with lysosomal adapter proteins and is degraded by the lysosome is consistent with the recent finding that sclerostin is found in extracellular vesicles positive for the lysosome associated protein LAMP1 (24).

**Fig. 3:**
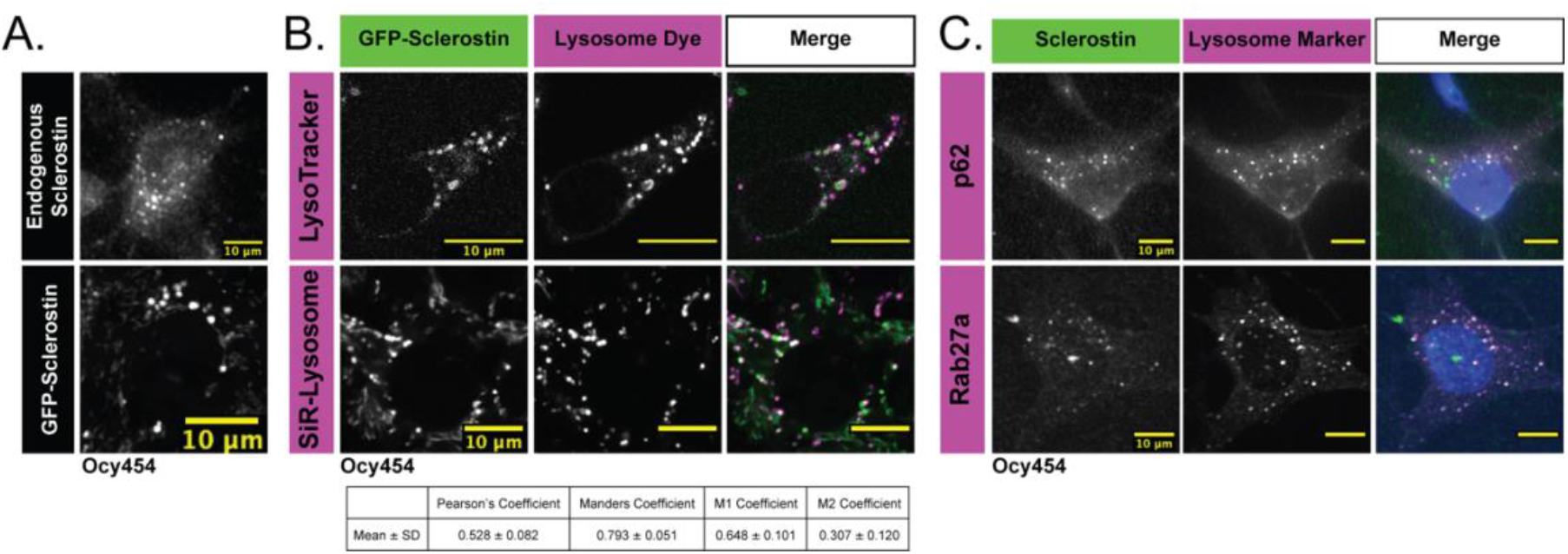
Endogenous and exogenous sclerostin co-localize with lysosomal markers. **(A)** Endogenous and GFP-tagged sclerostin both form discrete puncta in Ocy454 cells. **(B)** Ocy454 cells were transfected with GFP-sclerostin and lysosomes were visualized with Lysotracker (1mM, 1 hour) or siR-Lysosome (1μM, 4 hours). Z-stack images from GFP-sclerostin and siR-lysosome labeled cells were analyzed with the FIJI plug-in JaCOP for co-localization coefficients. M1 represents the overlap of sclerostin with siR-lysosome; M2 represents the overlap of siR-lysosome with sclerostin (n=3). Scale bar represents 10μm. **(C)** Ocy454 cells were fixed and stained for endogenous sclerostin and either p62 or Rab27a to evaluate co-localization.

Interestingly, sclerostin was also co-detected with Rab27a in Ocy454 cells (Fig. 3C). Rab27a is a small GTPase that is a master regulator of the trafficking, docking, and fusion of secretory lysosomes and also controls secretory granules in insulin-secreting beta-cells (25). This Rab27a-depedent secretory lysosome pathway also regulates RANKL secretion in osteoblasts as well (26), indicating a common mechanism in osteoblast-lineage cells for controlling the abundance of secreted proteins that control bone remodeling by diverting them to the lysosome.

### Lysosome activity is regulated by bone anabolic stimuli

Next, we examined if FSS or PTH altered lysosomal activity. Following a short, 1-minute bout of FSS and preceding sclerostin degradation, there is a transient increase in the autophagy cargo adaptor p62/sequestosome-1 protein abundance and the LC3 II/I ratio (Fig. 4A). Similarly, after 20 minutes of PTH exposure, p62/ sequestosome-1 is transiently increased prior to the loss of sclerostin protein, before returning to non-stimulated levels (Fig. 4B). Similarly, FSS induces an increase in lysosome activity, as determined using Magic Red Cathepsin-B activity assay (Fig. 4C). Together, these data support that osteocytes increase their lysosomal activity following bone anabolic stimuli and that this lysosomal activation precedes sclerostin degradation.

**Fig. 4.**
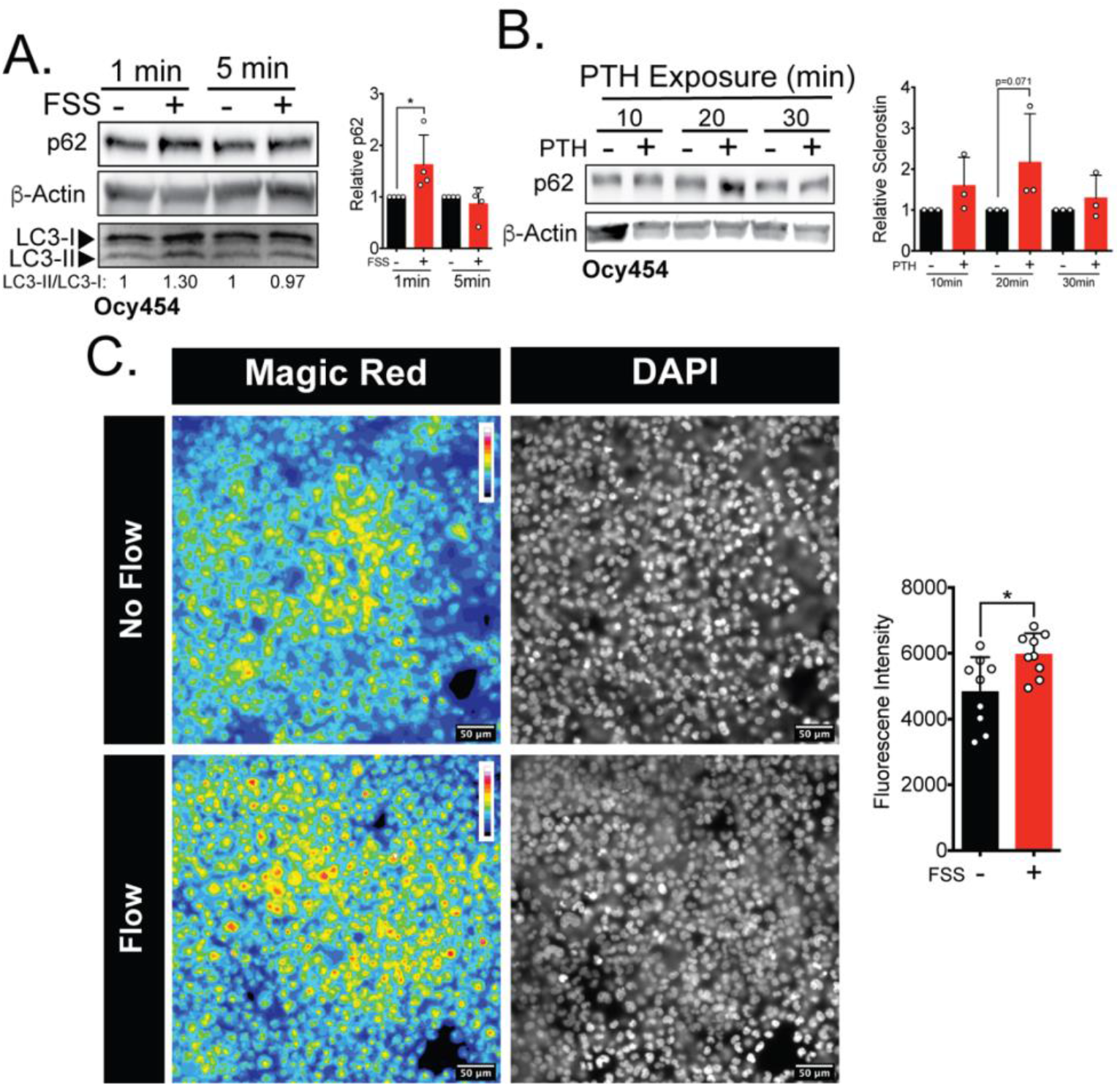
FSS and PTH treatment increase lysosomal activity. **(A)** Ocy454 cells were subjected to 1 minute or 5 minutes of FSS, lysed immediately post-flow, and western blotted for p62 and β-actin (n=4) and LC3 (n=1). **(B)** Ocy454 cells were treated with PTH (10nM) for the time indicated, lysed, and western blotted for p62 and β-actin (n=3). p62 abundance relative to β-actin was quantified. **(C)** UMR106 cells were subjected to FSS for 5 minutes, then Magic Red Cathepsin-B was applied for 10 minutes. Cells were then fixed and imaged for Magic Red intensity (n=9 independent images). Graphs depict mean ± SD. *p<0.05 by two-way ANOVA with Holm-Sidak post-hoc correction (A & B) or by unpaired two-tailed t-test (C).

### Nitric oxide is necessary and sufficient for rapid sclerostin degradation following FSS

To gain insight into the mechanisms regulating post-translational control of sclerostin degradation, we examined common regulators associated with both mechanical load and PTH responses. Both of these osteoanabolic cues activate CaMKII (15, 27, 28), and, using pharmacologic and genetic approaches, we previously reported that CaMKII activation is required for the rapid reduction of sclerostin by FSS in Ocy454 cells (15). Additionally, FSS and PTH regulate nitric oxide signaling (29, 30), and CaMKII regulates nitric oxide synthases (NOS) and vice versa (31, 32). Interestingly, both CaMKII and nitric oxide are implicated in protein degradation by the lysosome in other tissues (33-36). Despite nitric oxide being an established effector produced by bone cells in response to mechanical cues and PTH administration (29, 37-40), its direct biological consequence on the potent activation of osteoblasts and bone formation has remained incomplete. Here, we show for the first time that activation of nitric oxide signaling in Ocy454 cells with SNAP, a nitric oxide donor, induced a rapid decrease in sclerostin protein without increased CaMKII phosphorylation (Fig. 5A). When Ocy454 cells transfected with myc-sclerostin were treated with SNAP, a nitric oxide donor, in the presence of the lysosome inhibitor Bafilomycin A1, sclerostin degradation was prevented (Fig. 5B), confirming that nitric oxide is sufficient to drive the lysosomal degradation of sclerostin. Additionally, blocking nitric oxide production with L-NAME prevented the FSS-activated degradation of sclerostin protein at 5 minutes, as well as the FSS-induced increase in p62 at 1-minute post-FSS (Fig. 5C), supporting a role of nitric oxide in the activation of the lysosome and degradation of sclerostin. Additionally, Ocy454 cells treated with PTH in the presence of the CaMKII inhibitor, KN-93, failed to phosphorylate eNOS at serine 1177, suggesting that CaMKII activation may be upstream of NOS activation in this pathway (Fig. 5D). Combined, these results demonstrate that nitric oxide signaling following osteoanabolic cues is likely downstream of CaMKII and is both necessary and sufficient to induce lysosomal degradation of sclerostin protein. In total, these data establish that sclerostin is directed through a defined secretory pathway unless directed to the lysosome by mechano-transduction or PTH-initiated signaling.

**Fig. 5:**
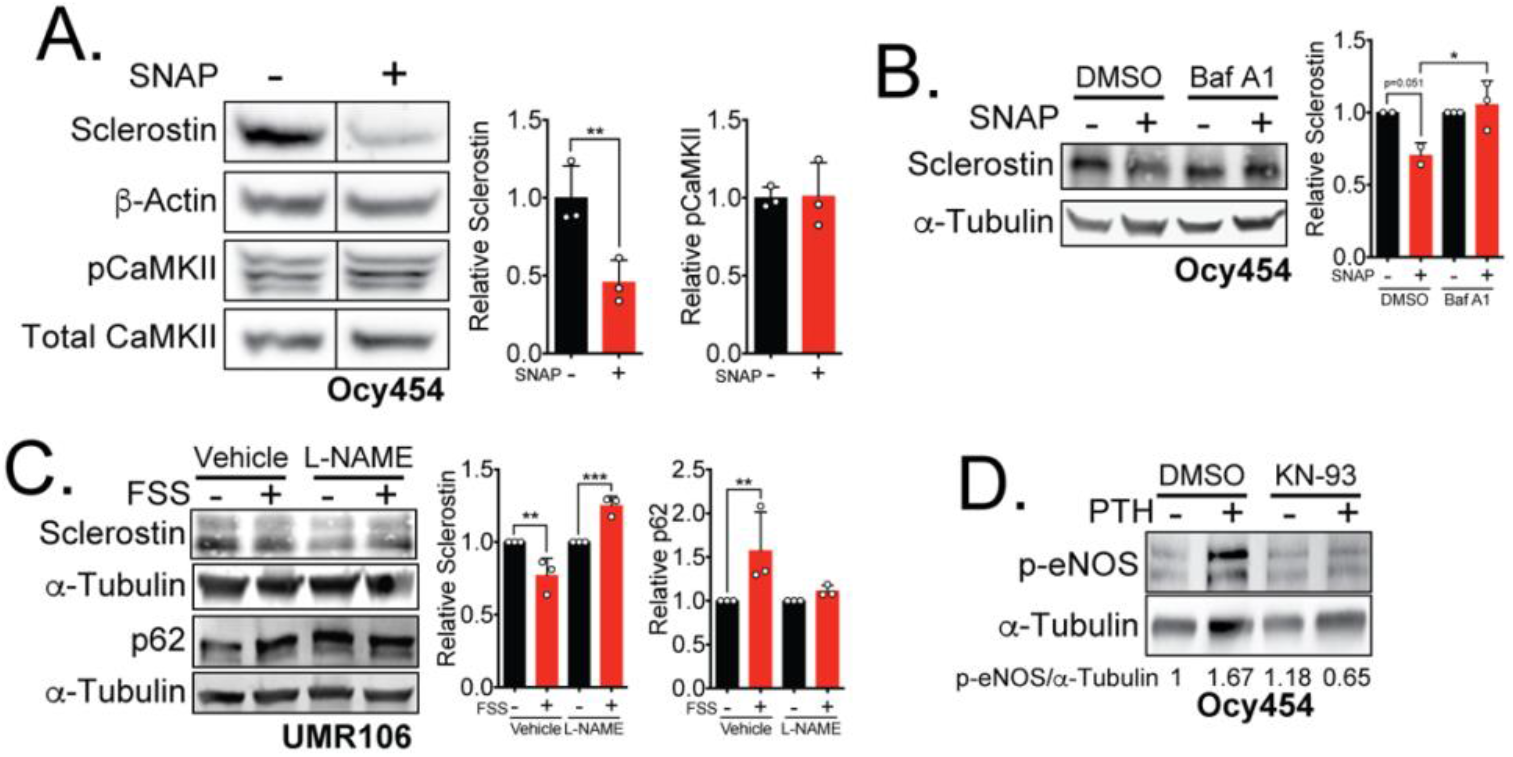
Nitric oxide contributes to the acute regulation of sclerostin protein downstream of CaMKII. **(A)** Ocy454 cells transfected with GFP-sclerostin were treated with vehicle (water) or 10μM SNAP and lysed after 5 minutes. Western blots were probed for sclerostin, α-tubulin, pCaMKII, and total CaMKII (n=3). For each antibody, blots are from a single gel and exposure; a vertical black line indicates removal of irrelevant lanes. **(B)** Ocy454 cells transfected with myc-tagged sclerostin were treated with DMSO or Bafilomycin A1 (100nM) for 30 minutes, then treated with SNAP for 5 minutes and lysed. Western blots were probed for sclerostin and α-tubulin (n=2-3). **(C)** UMR106 cells were treated with vehicle or L-NAME (1mM) for 1 hour then exposed to 1 or 5 minutes of FSS. Lysates from cells exposed to 1 minute of FSS were probed for p62 and α-tubulin abundance and lysates from cells exposed to 5 minutes of sclerostin were probed for sclerostin and α-tubulin abundance (n=3). (**D**) Ocy454 cells were treated with DMSO or KN-93 (10μM) for 1 hour. PTH (10nM) was then added for an additional 10 minutes, then lysed. Western blots were probed for p-eNOS and α-tubulin (n=1). Graphs depict mean ± SD. *p<0.05, **p<0.01, ***p<0.001. by unpaired two-tailed t-test (A) or two-way ANOVA with Holm Sidak post-hoc test (B & C).

### Lysosomal function and the pathway regulating sclerostin degradation are necessary for the *in vivo* bone mechano-response

To translate our findings to *in vivo* models, we utilized two complimentary approaches. First, we tested effects of transiently disrupting the lysosome, and thus sclerostin degradation, just prior to mechanical loading and quantified the subsequent effects on load induced bone formation. Second, we acutely targeted an essential early step in the mechano-signaling pathway (NOX2 activation) that leads to sclerostin degradation just prior to each bout of loading and then examined the subsequent mechanically induced bone formation. To determine the contribution of lysosomal degradation to load-induced bone formation *in vivo*, we performed ulnar loading in mice treated with vehicle or the lysosome inhibitor Bafilomycin A1 once a day for four consecutive days. After the final day of loading, the bone surfaces were labeled with Alizarin Red and Calcein and dynamic histomorphometry was performed to assess *de novo* bone formation. Importantly, Bafilomycin A1 was not administered during the inter-label period when bone formation was monitored. Acute administration of Bafilomycin A1 4 hours prior to each bout of ulnar load blocked the subsequent load-induced increase in periosteal bone formation rate (Ps.BFR) (Fig. 6A and 6B) and mineral apposition rate (Ps.MAR) (Fig. 6A and 6C) observed 14 days after the initiation of the experiment. In contrast, the administration of Bafilomycin during the first four days of the experiment had no effect on the Ps.BFR or Ps.MAR in the contralateral non-loaded limb.

**Fig. 6:**
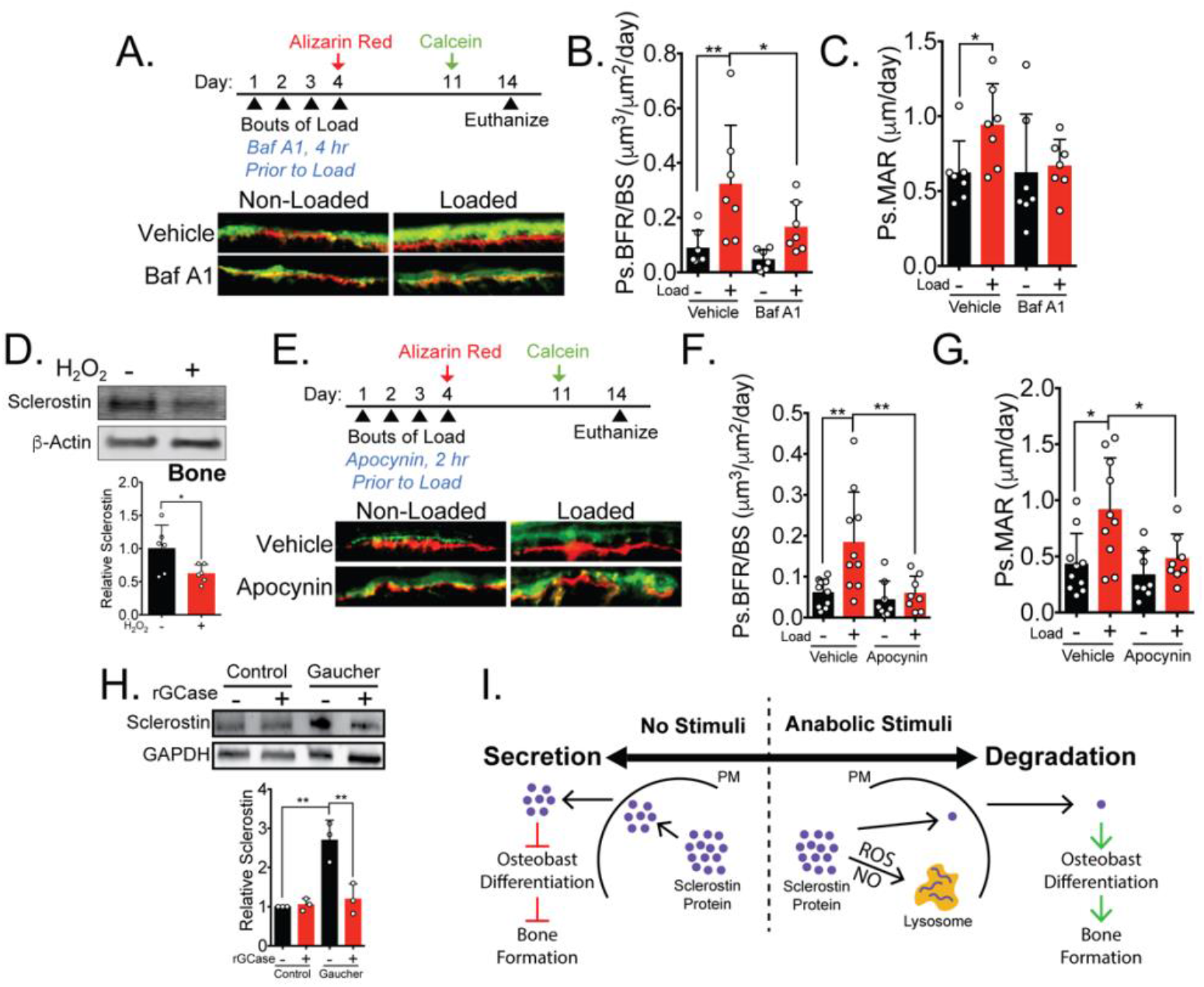
Lysosome function and the pathway upstream of sclerostin degradation are necessary for the in vivo mechano-response. **(A)** 15-week old male C57Bl/6 mice treated with vehicle (saline + 4% DMSO, n=7 independent animals) or Bafilomycin A1 (1mg/kg, n=7 independent animals) were forearm loaded (3.2N, 90sec, 2Hz) and labeled with calcein and alizarin red at the indicated times for dynamic histomorphometry. Representative periosteal double labeling are shown. **(B)** Periosteal bone formation rate (Ps.BFR) and **(C)** periosteal mineral apposition rate (Ps.MAR) were calculated. **(D)** Dissected cortical osteocyte-enriched ulnae and radii were treated with 100uM hydrogen peroxide *ex vivo* for 5 minutes before homogenization. Western blots probed for sclerostin and β-actin (n=6 independent animals). Sclerostin abundance relative to β-actin was quantified. **(E)** 13-week old male C57Bl/6 mice treated with vehicle (saline, n=10 independent animals) or Apocynin (3 mg/kg, n=8 independent animals) were forearm loaded (3.4N, 90sec, 2Hz) and labeled with calcein and alizarin red at the indicated times for dynamic histomorphometry. Representative periosteal double labeling are shown. **(F)** Periosteal bone formation rate (Ps.BFR) and **(G)** periosteal mineral apposition rate (Ps.MAR) were calculated. **(H)** Human iPSCs-derived osteoblasts from either control (non-diseased) or Gaucher disease patients were treated with vehicle or recombinant glucocerebrosidase (rGCase, 0.24U/mL) for 5 days, then lysed for western blotting. Western blots were probed for sclerostin and GAPDH (n=3 independent patient-derived iPSC lines/group). Graphs depict mean ± SD. *p<0.05, **p<0.01 by two-way ANOVA with Holm-Sidak post-hoc correction. **(I)** Osteoanabolic stimuli, working through reactive oxygen (ROS) and reactive nitrogen species (RNS), direct sclerostin to the lysosome for degradation. This results in reduced sclerostin to allow for bone formation. PM: Plasma membrane; ROS: Reactive Oxygen Species; NO: Nitric Oxide.

Next, we targeted the upstream pathway implicated in the degradation of sclerostin. Our prior work revealed that NOX2-derived ROS is an essential early step in the mechanotransduction pathway converging on sclerostin protein loss (15). We demonstrated that hydrogen peroxide (15) or ROS generated by a genetically encoded photoactivatable protein, KillerRed, was sufficient to activate CaMKII and decrease sclerostin protein abundance in Ocy454 and UMR106 cells, respectively (Supplemental Fig. 3A and 3B). To understand the direct contribution of ROS to sclerostin regulation in an intact bone, isolated ulnae and radii were treated with hydrogen peroxide for 5 minutes *ex vivo* to asses sclerostin abundance. Similar to Ocy454 cells treated with hydrogen peroxide *in vitro* (15), sclerostin abundance was significantly reduced in bones treated with hydrogen peroxide (Figure 6D). To link the mechano-activated pathway regulating of sclerostin protein to bone formation *in vivo*, we targeted NOX2 activity with Apocynin *in vivo* in the context of ulnar loading. Acute administration of Apocynin 2 hours prior each bout of ulnar loading blocked the subsequent load-induced increase in periosteal bone formation rate (Ps.BFR) (Fig. 6E and 6F) and mineral apposition rate (Ps.MAR) (Fig. 6E and 6G) observed 14 days after the initiation of the experiment. As with Bafilomycin A1, Apocynin was not administered during the inter-label period when bone formation was monitored. Importantly, the administration of Apocynin during the first four days of the experiment had no effect on the subsequent Ps.BFR or Ps.MAR in the contralateral non-loaded limb.

When interpreted together, these results demonstrate that blocking lysosomal function or the mechano-pathway controlling sclerostin degradation both yielded a predicted inhibition of load-induced bone formation. These data strongly support the relevance of post-translational control of sclerostin to skeletal physiology and adaptation to mechanical loading.

### Disrupted lysosomal function in Gaucher Disease leads to sclerostin dysregulation

Having defined sclerostin regulation by the lysosome, we probed the clinical relevance using induced pluripotent stem cell (iPSC)-derived osteoblasts from Gaucher disease, a lysosomal storage disorder in which patients lack the lysosomal hydrolase glucocerbrosidase (GCase). As with many lysosomal storage disorders, patients with Gaucher disease exhibit skeletal dysplasias and low bone mass (41). Gaucher disease iPSC-derived osteoblasts display defective osteoblast differentiation and mineralization caused by defects in Wnt/β-catenin signaling, an effect that is reversed by treatment with recombinant GCase (42). Accordingly, we speculated that this suppression of β-catenin signaling and osteoblast differentiation and function may be a consequence of defects in sclerostin control, as sclerostin inhibits both of these processes. Consistent with lysosomal degradation of sclerostin, we observed that iPSC-derived osteoblasts from Gaucher patients had significantly increased levels of sclerostin compared to iPSC-derived osteoblasts from healthy patients without Gaucher disease (Fig. 6H). Further, treating Gaucher iPSCs-derived osteoblasts with recombinant GCase, which restores lysosomal function and osteoblast differentiation (42), restored sclerostin abundance to control levels (Fig. 6H). Since sclerostin acts as an inhibitor of Wnt/β-catenin signaling, a phenotype-driving defect in these cells (42), these data expand the mechanistic basis for the observed defects in Gaucher-derived osteoblasts and suggest sclerostin as a therapeutic target in for bone loss in Gaucher disease.

## Discussion

Our data show the unexpected finding that osteocytes respond to distinct osteoanabolic cues, mechanical load and PTH, by redirecting sclerostin from a secretory pathway to the lysosome for rapid degradation (Fig. 6I). This signaling event is mediated by activation of ROS and nitric oxide signaling, an increase in lysosomal activity, and the post-translational degradation of sclerostin by the lysosome. Further, we demonstrate that both lysosomal activity and activation of the upstream of mechano-signaling pathway are required for mechanically induced bone formation *in vivo*. Importantly, these findings inform a new model of osteocyte mechano-transduction and regulation of sclerostin-containing secretory lysosomes that integrates many of the molecular effectors of the osteocyte mechano-response, including calcium, the cytoskeleton, nitric oxide, and sclerostin (17-20) and unifies many important discoveries related to osteoanabolic signals that act through sclerostin. These findings also describe a role of nitric oxide production, a canonical response to mechanical load in bone of previously unclear functional consequence, in the regulated degradation of sclerostin protein. Finally, we link sclerostin degradation not only to skeletal physiology in mice but also to human disease using Gaucher disease iPSCs.

These data do not preclude a transcriptional control of the *Sost* gene by osteoanabolic stimuli (43, 44). Rather, they add an important downstream check point for regulating sclerostin protein bioavailability with remarkable temporal control. It may be that post-translational control is the first line temporal response to a bone anabolic stimulus, whereas long term stimulation leads to genome level transcriptional control of the *Sost* gene. Indeed, longer periods of loading are required to observe *Sost* mRNA decreases than are required for loss of protein described here. The transient regulation of sclerostin protein abundance may be important for controlling the local response to mechanical stress, as bone formation is asymmetrical across the tissue (45-47). We speculate that the rapid and transient nature of sclerostin degradation may be critical to the precise anatomical positioning of new bone formation following an anabolic stimulus. First, the transient nature of sclerostin degradation ensures that deficits in local sclerostin caused by osteocytes experiencing strain are unlikely to affect distal sites where mechanical cues are absent. Thus, *de novo* bone formation will be local. Second, given that a single load event may have minimal impact on the local sclerostin abundance and the de-repression of canonical Wnt/β-catenin signaling, it may be the integral of load over time (or activation of PTH receptor signaling over time) that determines the magnitude of the biological consequence of a bone anabolic agent, as sustained local activation of sclerostin degradation may lead to a more robust osteoanabolic response.

In isolation, the effects of inhibiting either NOX2-dependent ROS or lysosomal function on load-induced bone formation could be influenced by the broad pharmacological impact of these drugs. However, that two distinct pharmacological inhibitors targeting different steps in the pathway controlling sclerostin protein abundance yielded a similar effect on bone formation strongly supports our conclusions and is consistent with our *in vitro* findings. This conclusion is strengthened by two other important considerations: firstly, there was also no statistical effect of either Apocynin or Bafilomycin A1 on bone formation rate or mineral apposition rate in the contralateral, non-loaded limb, and, secondly, the inhibitors were only present for each bout of ulnar load, but were absent during the seven-day bone forming period, indicating that disruption of the acute downstream response to the mechanical cue, not the ability of the osteoblasts to build bone, caused the reductions in bone formation.

The identification of both exogenous and endogenous sclerostin in discrete pucta associated with Rab27a was surprising. Rab27a is associated with secretion through a specialized secretory lysosome pathway (48). Notably, this Rab27a-associated secretory lysosome pathway functions in osteoblasts to control RANKL secretion (26). Whether this mechano-pathway and regulated degradation extends to osteocyte derived RANKL was not analyzed in the present study, but may be physiologically significant as osteocytes are the primary source of bioactive RANKL directing bone resorption in response to physiologic cues (49, 50). In support of the notion that sclerostin is packaged into secretory lysosomes, our *in silico* analysis found the conservation of lysosomal signal sequences, as described above, in addition to the presence of a secretory signal peptide at residues 1-23 (Supplemental Fig. 2).

Our data support that sclerostin is directed through a defined secretory pathway for exocytosis, then it is either secreted (no stimulus present) or trafficked to the lysosome for degradation (recent mechanical stress or PTH exposure). The rapid, controlled degradation of osteocyte sclerostin has intriguing parallels to crinophagy, a specialized autophagic process in peptide-secreting cells in which cellular cues elicit the re-routing of secretory proteins to the lysosome (51). Crinophagy is well described in pancreatic beta-cells, in which insulin-filled vesicles that are typically targeted for secretion are instead shuttled to the lysosome for degradation (36). In beta-cells, crinophagy is a nitric oxide-regulated process, with dysfunctional nitric oxide synthase (NOS) leading to increases in insulin-containing secretory granules that are unable to properly fuse with lysosomes (36). Additionally, Rab27a is associated with the secretion of insulin-filled granules in pancreatic beta-cells (25). While mechano-activation of crinophagy has not been reported previously, it is intriguing to consider that mechano-activated crinophagy may be more broadly applicable. Indeed, while these data are focused on bone, our initial report on this mechano-transduction pathway in osteocytes (15) and this report are mechanistic extensions of the pathway our team first discovered in heart (52) then in skeletal muscle (53, 54).

Using Gaucher disease as a model system, this work links lysosomal function to sclerostin regulation in disease. Gaucher patients experience many skeletal dysplasias, including low bone mass and osteoporosis. Though this cell autonomous defect in Gaucher osteoblasts has previously been linked to dysregulated β-catenin that impaired osteogenic differentiation and mineralization capacity (42), the present findings reveals a possible mechanism by which β-catenin is suppressed. Sclerostin is a canonical Wnt/β-catenin antagonist; therefore, the increased sclerostin levels observed here in Gaucher disease iPSCs are likely to lead to the previously reported decrease in β-catenin activation and osteoblast differentiation (42). Likewise, we showed that restoring expression of the missing hydrolase, glucocerebrosidase, in Gaucher disease iPSCs lowers sclerostin abundance, which could explain the rescue in β-catenin activation and osteoblast differentiation reported previously (42). Our data support the exploration of therapeutics that target sclerostin to treat low bone mass symptoms in patients with lysosomal storage disorders.

That sclerostin protein abundance is post translationally controlled may have important implications in age-related osteoporosis and the reduced sensitivity of osteocytes to mechanical cues with aging (55, 56). Lysosome activity is diminished with age (57, 58), including in the osteocyte during age-related bone loss (59), an effect that could impair sclerostin degradation and new bone formation even in the face of osteoanabolic signals. While undoubtedly multifaceted in its impacts, targeting autophagy and lysosome activity to improve bone mass in aging has been proposed (60, 61), and these data suggest that impacts on sclerostin bioavailability might contribute mechanistically to its efficacy.

Together, these discoveries provide key insights into the unexpected, rapid regulation of osteocyte sclerostin protein by the lysosome and reveal new therapeutic targets that can be exploited to improve bone mass in conditions such as osteoporosis. Additionally, targeting sclerostin may be important to interventions to improve bone mass in lysosomal storage disorders, like Gaucher disease.

## Materials and Methods

### Chemicals and Reagents

Bafilomycin A1 (*in vitro* studies, #54645), MG-132 (#2194), Brefeldin-A (#9972) Cycloheximide (#2112), and antibodies against Thr 286 pCaMKII (#12716S), total CaMKII (3362S), p-eNOS (Ser 1177, #9571), p62 (#23214), LC3B (#2775), Rab27a (#69295), and Prolong Gold Antifade Reagent with DAPI (#8961S) were from Cell Signaling Technologies. REVERT® Total Protein Stain (827-15733) was from Licor. Anti-sclerostin antibodies (#AF1589) were purchased from R&D Systems. Bafilomycin A1 (*in vivo* studies, #88899-55-2) was from Research Products International. Leupeptin (#EI8), N_ω_-Nitro-L-arginine methyl ester hydrochloride (L-NAME, N5751), and GAPDH (MAB374) were from Millipore. Parathyroid Hormone (1-34, #P3109-24D) (PTH) was from US Biological Life Sciences. Alizarin Red (#A3882), Calcein (#C0875), Apocynin (178385), S-Nitroso-N-acetyl-DL-penicillamine (SNAP, N3398), and antibodies against β-Actin (A1978), α-Tubulin (T9026), Pro-Collagen Type I, A1 (Col1α1) (ABT257), and Magic Red Cathepsin B Detection Assay Kit (CS0370) were from Sigma. siR-Lysosome (CY-SC016) was from Spirochrome. KN-93 (202199) was from Santa Cruz. GFP-tagged human sclerostin (#RG217648)- and myc-tagged mouse sclerostin (#MR222588) were purchased from Origene. KillerRed plasmid (FP966) was purchased from Evrogen. Recombinant human GCase (rGCase) (Cerezyme®) was obtained from patient infusion remnants. 2’,7’-dichlorofluorescein (DCF, D399) was purchased from Invitrogen. Halt™ Protease and Phosphatase Inhibitor Cocktail (EDTA-free) Lysotracker (L7528), SuperBlockPBS (37515), Donkey anti-Goat IgG Alexa Fluor 546 (A-11056), and Chicken anti-Rabbit Alexa Fluor 488 (A-21441) were from Thermo Fisher Scientific. JetPrime Transfection kit was from PolyPlus Transfection. Modified RIPA lysis buffer contained 50 mMTris-HCl pH 8.0, 150 mM NaCl, 1.0% NP-40, 0.5% sodium deoxycholate, 0.1% SDS, 10 mM Na_4_P_2_O_7_, 10 mM 2-glycerolphosphate, 10 mM NaF, 10 mM EDTA, 1 mM EGTA, 1× HALT phosphatase and protease inhibitor cocktail.

### Cell Culture

UMR106 cells (purchased from ATCC, CRL-1661) were cultured in Dulbecco’s modified essential medium (DMEM) supplemented with 10% fetal bovine serum (FBS), and maintained at 37°C and 5% CO2, as described (62). Ocy454 cells (provided by P. Divieti-Pajevic, Boston University) were cultured on type I rat-tail collagen coated plates in α-minimal essential medium (αMEM) supplemented with 10% FBS and maintained at 33°C and 5% CO_2_ (13, 14). Prior to experimentation, cells were seeded into tissue-culture-treated vessels and maintained at 37°C and 5% CO_2_. The iPSC from a patient with type 2 Gaucher disease and a control subject used in this study have been previously described (42). Their genotypes are: W184R/D409H and WT/WT (Control MJ). Control and Gaucher disease iPSC were differentiated to osteoblasts as described (42). Briefly, embryoid bodies from WT and Gaucher iPSCs were transferred to 0.1% (w/v) gelatin-coated plates and cultured in MSC medium [high glucose DMEM (Invitrogen), 20% FBS (Hyclone), 1 mM l-glutamine and 100 U/ml Pen/Strep (Invitrogen)] to generate mesenchymal stem cells (MSCs). To generate osteoblasts, the WT and Gaucher MSCs were plated at a density of 2 ×10^4^ cells/cm^2^ and were cultured in osteoblast differentiation media [MSC media supplemented with 10 mM beta-glycerophosphate (Sigma), 100 µM dexamethasone (Sigma) and 50 μg/ml ascorbic acid (Sigma)] for 3-4 weeks, as described (42). Recombinant glucocerebrosidase (rGCase, 0.24 U/ml) was added to the cultures with each media change (42).

### Fluid Flow

Ocy454 and UMR106 cells were exposed to fluid flow using a custom FSS device (15, 63). Media was removed, cells were rinsed in a Hepes-buffered Ringer solution containing 10 mM Hepes (pH 7.3), 140mM NaCl, 4mM KCl, 1mM MgSO_4_, 5mM NaHCO_3_, 10 mM glucose, and 1.8 mM CaCl_2_. Ringer solution was also used as fluid flow buffer. Cells were exposed to 1-5 minutes of FSS (4 dynes/cm^2^), as indicated, and lysed in a modified RIPA buffer plus HALT protease and phosphatase inhibitors at the time indicated in each experiment.

### Cell Treatments

To block cellular degradation pathways, cells were pre-treated for 30 minutes or 4 hours with Bafilomycin A1 (100nM), Brefeldin A (2μM), MG-132 (10μM), Leupeptin (200μM, 6 hours), or DMSO (0.1%) as the vehicle control in Ringer solution, as indicated. For PTH treatment, cells were treated with PTH (1-34) (10nM) or Ringer solution as the vehicle for up to 30 minutes. For SNAP treatment, cells remained in the media they were plated in and SNAP dissolved in sterile water was added to a final concentration of 10μM. Vehicle was sterile water. To block the lysosomal function before SNAP treatment, cells were treated with Bafilomycin A1 (100nM) for 30 minutes prior to the addition of SNAP. Cells were lysed 5 minutes after addition of SNAP. For L-NAME treatment, L-NAME was dissolved in sterile water, then diluted into Ringer solution (1mM). UMR106 cells were pre-treated with L-NAME or vehicle control 1 hour prior to FSS, then exposed to 5 minutes of FSS, and lysed immediately after. Ocy454 cells were treated with 10μM KN-93 for 1 hour prior to the addition of PTH (10nM) for an additional 10 minutes.

### Sequence Alignment

Amino acid sequences were acquired from the National Center for Biotechnology Information’s (NCBI) protein database. Accession numbers are as follows: mouse (AAK13455); human (AAK16158.1); rat (EDM06161.1); cow (NP_001159986.1); chicken (XP_024999845.1). Sequence alignment was done using NCBI’s Constraint-based Multiple Alignment Tool (COBALT). Lysosomal signal sequences (23) and the secretory signal peptide were annotated manually.

### Degradation Assays

Ocy454 cells were treated with Cycloheximide (CHX, 150µg/mL) and either DMSO (0.1%), Bafilomycin A1 (100nM), Brefeldin A (2µM), or MG-132 (10µM) diluted in supplemented αMEM for 4 hours or with Leupeptin 200µM for 6 hours at 37°C and 5% CO_2_. Cells were then exposed to 5 minutes of FSS as described above in Ringer containing the appropriate treatment with CHX. Cells were lysed in a modified RIPA buffer + HALT protease and phosphatase inhibitors and collected for western blotting at 5- and 30-minutes post-flow. For basal degradation assays, UMR106 cells were treated with 150µg/mL CHX for 0, 1, 2, or 4 hours and lysed. A best fit linear line was constrained through y=1 to determine protein half-life.

### Transient Transfections

Ocy454 cells were seeded in 96-well plates at a density of 20,000 cells/well and incubated at 37°C for 24 hours. Transient transfections were performed using 0.025 µg/well GFP-tagged sclerostin or 0.05 µg/well Myc-tagged sclerostin DNA mixed with 5uL/well JetPrime buffer and 0.1 µl/well JetPrime reagent, as described (15). After 16 hours of incubation at 37°C, the transfection medium was replaced with complete αMEM. The cells were incubated an additional 24 hours at 37°C prior to experiments. UMR106 cells for KillerRed experiments were plated on glass bottom 10mm dishes at a density of 50,000 cells/dish. Transient transfections were performed using 2 µg/well KillerRed and 4 or 8 µl/well Jetprime reagent. 16 hours after incubation, transfection medium was replaced with complete DMEM and were incubated an additional 37°C prior to experiments.

### Fluorescence Co-Localization

Ocy454 cells were seeded at 10,000 cells/well and grown on a glass-bottom 96-well plate. Cells were transfected with GFP-Sclerostin as described above, where indicated. For immunofluorescence staining, cells were fixed with 1% paraformaldehyde, permeabilized with 0.1% Triton-X in PBS, and blocked with SuperBlockPBS for 1 hour. Primaries diluted in SuperBlockPBS against p62, Rab27a, and sclerostin were used at 1:250 and incubated overnight at 4C. Chicken anti-Rabbit 488 and Donkey anti-Goat 546 diluted in SuperBlockPBS were used at 1:100 and incubated for 3 hours at room temperature. Cells were mounted with ProLong Gold Antifade with DAPI and imaged with a Nikon Ti2 microscope with a SpectraX Light Engine and a ds-Qi2 Monochrome camera. For live cell imaging, Ocy454 cells were transfected with GFP-Sclerostin as described above. Lysosomes were labeled Lysotracker (1μM) for 1 hour, then Nikon C2 confocal microscope. For co-localization quantification, Ocy454 cells transfected with GFP-sclerostin were treated with siR-Lysosome (1μM) for 4.5 hours at 37°C to label lysosomes. Cells were imaged on a Nikon C2 confocal microscope and Z-stacks with 0.5μm steps were obtained in each well. These Z-stacks were then denoised using Nikon Denoise aI (NIS Elements 5.2). Co-localization in the denoised Z-stacks using the Mander’s Coefficients was measured using the FIJI plugin, JaCOP (64).

### Magic Red Cathepsin B Activity Assay

UMR106 cells were exposed to FSS at 4 dynes/cm^2^ for 5 minutes. Magic Red and Hoechst (1x) reagent were then was added to all wells for 10 minutes. Cells were washed three times with warm 1X PBS and then fixed for 10 minutes with 1% paraformaldehyde. Cells were washed one time with 1X PBS and imaged at 20x with a Nikon Ti2 microscope with a SpectraX Light Engine and a ds-Qi2 Monochrome camera. Average Magic Red intensity was measured by subtracting a mask image of the nuclei from the Magic Red signal. Magic Red signal was then measured on a per pixel basis and average intensities were calculated.

### KillerRed Imaging and Protein Isolation

For imaging, UMR106 cells transfected with KillerRed were loaded with DCF (10μM, 30 min, 37°C) diluted in Ringer solution to track ROS production. After loading, cells were washed with fresh Ringer solution. Cells were stimulated with LED light for 40 seconds. DCF and KillerRed signals were imaged before and after exposure to LED light in the same cells. For western blotting, cells transfected with KillerRed were stimulated with LED light for 5 minutes and were lysed 5 minutes after the end of light exposure. No light controls were treated the same but were not exposed to light.

### Animals

Male and female, age matched C57BL/6 mice were purchased from Jackson Laboratory. Mice were group housed in micro-isolator cages, and food (standard rodent chow) and water were available *ad libitum*. Mice were maintained on a 12-h-light–12-h-dark cycle. Experiments were conducted on 13-16 week old mice, as indicated. All animal protocols were approved by the Animal care and Use Committee at the University of Maryland School of Medicine.

### *Ex vivo* Treatments

Ulnae and radii were dissected from surrounding soft tissues, epiphyses were cut, and marrow was flushed with Ringers solution. Bones were acclimated in complete αMEM at 37°C and 5% CO_2_ for at least 20 minutes before moving them into Ringers solution with vehicle control or 100μM hydrogen peroxide for 5 minutes at 37°C. Bones were then removed from the Ringers solution and were homogenized in RIPA buffer + HALT protease and phosphatase inhibitor cocktail using a Bullet Blender (Next Advance), as described (65, 66). Extracts were subsequently used for western blotting analysis, as described below.

### *In vivo* Loading

For acute isolation of protein, 16-week old female C57BL/6 mice were subjected to a single, acute bout of ulnar loading. Briefly, mice were anesthetized (isoflurane) and their left upper limb was placed in a horizontal orientation in a uniaxial load device (Aurora Scientific, 305C-FP). A small pre-load was applied (0.4N) and then the forearm was cyclically loaded with a sinusoidal wave at 2Hz and a peak force of 2.8N for 90sec, as described (67, 68). Five minutes following load, loaded and contralateral non-loaded (control) ulnae and radii were dissected from surrounding soft tissue, epiphyses removed, flushed of marrow, and homogenized in RIPA buffer + HALT protease and phosphatase inhibitor cocktail using a Bullet Blender (Next Advance), as described (65, 66). Extracts were subsequently used for western blotting analysis, as described below.

For dynamic histomorphometry following mechanical stimulation, 13-week (Apocynin experiments) or 15-week old (Bafilomycin A1 experiments) male C57BL/6 mice were subjected to four consecutive days (days 1 – 4) of forearm loading at 3.4N (Apocynin experiments) or 3.2N (Bafilomycin A1 experiments) at 2Hz for 90sec, as described above. For Apocynin experiments, Apocynin (3 mg/kg dissolved in saline) or vehicle control (saline) were injected 2 hours prior to each bout of ulnar loading. The contralateral limb served as a non-loaded control in all loading experiments. Following loading, mice were returned to their cages for unrestricted activity. For Bafilomycin A1 experiments, Bafilomycin A1 (1 mg/kg, i.p., 4% DMSO in saline) or vehicle control (saline with 4% DMSO) was injected 24 hours before the first bout of loading (day 0) and 2 hours prior to each bout of loading. For both inhibitors, treatment occurred only during the loading phase and not during the subsequent monitoring of bone formation by dynamic histomorphometry.

To assess cortical bone formation rate during the post-load period, animals were injected with Alizarin Red (30mg/kg, i.p.) after the final bout of loading (day 4) and injected with Calcein (30mg/kg, i.p.) on day 11. Three days later (day 14), animals were euthanized, loaded and contralateral non-loaded ulnae and radii were dissected from surrounding soft tissue and stored in 100% ethanol. Bones were then placed in 30% sucrose in PBS at 4°C overnight. Bones were embedded in optimal cutting temperature compound (OCT) and sectioned at 5µm thickness using the Kawamoto Film Method (69). Three sections were collected for each ulna and averaged to obtain a single value for each parameter in each animal. Fluorescent labels were visualized with a Nikon Ti2 microscope at 20x using a Nikon Ri2 monochrome camera and cortical bone parameters were quantified using BioQuant 2019 Software. Parameters included: periosteal mineral apposition rate (Ps.MAR) and periosteal bone formation rate (Ps.BFR), as defined by the ASBMR nomenclature guidelines (70). Two animals from the Apocynin treated group, three animals from the 4% DMSO in saline vehicle group, and three animals from the Bafilomycin A1 group were excluded from analysis due to lack of periosteal labeling from one of the fluors, precluding calculation of meaningful measures of dynamic histomorphometry.

### Western Blotting

Western blotting of whole cell extracts following appropriate treatments or extracts isolated from murine long bone was done as previously described (66). Briefly, equal amounts of protein were loaded on SDS-PAGE gels, electrophoresed, and transferred to PVDF membranes. Membranes were blocked in 5% nonfat dry milk and 3% BSA in phosphate buffered saline with 0.1% Tween-20 for all sclerostin blots. All other blots were blocked in 5% nonfat dry milk in phosphate buffered saline with 0.1% Tween-20. Primary antibodies dilutions were: Sclerostin (1:500); GAPDH (1:2500); active β-Catenin (1:1000); β-Actin (1:5000), α-Tubulin (1:2000), p-eNOS (1:1000); pro-collagen type 1 (Col1a1) (1:1000), pCaMKII (1:1000), total CaMKII (1:1000), p62 (1:500), and LC3B (1:500). Antibodies were detected using horseradish peroxidase-conjugated secondary antibody (1:1000-5000) (Cell Signaling Technology) and visualized with enhanced chemiluminescence reagent (Bio-Rad) and analyzed with ImageLab software (Bio-Rad) or fluorescent Licor secondary antibodies (1:20,000) were used and visualized with Licor Oddysey CLx and analyzed using Image Studio v5.

### Statistical Analysis

Experiments were repeated a minimum of three times with triplicate samples unless indicated otherwise. For dynamic histomorphometry experiments, animals were randomized into groups by weight. Mice were assigned a random number, and the experimenters were blinded to animal treatment and experimental group during collection and analysis period. Graphs show means, with error bars indicating SD or SEM, as indicated. All statistical test applied to experimental data were carried out in GraphPad Prism 8.0. Data were compared with two-tailed unpaired t-tests or with a two-way ANOVA with Holm-Sidak post-hoc correction, as indicated. A p value of <0.05 was used as a threshold for statistical significance.

## Abbreviations

Asn: Asparagine
Baf A1: Bafilomycin A1
Bref A: Brefeldin A
Ca^2+^: Calcium
CaMKII: Calcium/Calmodulin-Dependent Kinase II
FDA: Food and Drug Administration
FSS: Fluid Shear Stress
GCase: Glucocerebrosidase
GFP: Green Fluorescent Protein
GlcNAC: N-Acetylglucosamine
iPSC: Induced Pluripotent Stem Cell
NOX2: NADPH Oxidase 2
Ps.BFR: Periosteal Bone Formation Rate
Ps.MAR: Periosteal Mineral Apposition Rate
PTH: Parathyroid Hormone
PTHrP: Parathyroid Hormone Related Peptide
RANKL: Receptor activator of nuclear factor kappa-B ligand
ROS: Reactive Oxygen Species
Ser: Serine
Thr: Threonine
TRPV4: Transient Receptor Potential Vanilloid 4
Tyr: Tyrosine
Φ: Hydrophobic Amino Acid Residue

## Acknowledgments

The Ocy454 cells were provided by P. Divieti-Pajevic (Boston University) through support from the Center for Skeletal Research Core at Massachusetts General (NIH P30 AR075042).

## Author Contributions

NRG, KMW, JSL, RJK, CWW and JPS designed the study; NRG, KMW, HCJ, OMT, JSL, JML, MPS, MH, RJK, CWW and JPS performed research and analyzed data; NRG, RJK, CWW, and JPS wrote the paper; NRG, RJK, CWW, and JPS prepared the figures; RJK, CWW, and JPS supervised the research; RAF, CWW, and JPS acquired funding to support the research. All authors reviewed and approved the final manuscript.

## Funding

This work was supported funding from NIH (AR071614, JPS and CWW; AR071618, HL142290, CWW; GM008181, NRG and JSL; and AR007592 KMW); Maryland Stem Cell Research Fund (2018-MSCRFD-4246, RAF); the Children’s Gaucher Research Fund (RAF); and the AHA (#19POST34450156, HCJ).

## Competing Interests

Declaration of competing interest JSL, CWW, and JPS hold two patents related to this work. One for the custom fluid shear device used for these experiments (US Patent No US 2017/0276666 A1) and a second for the targeting microtubules (part of this mechano-transduction pathway) to improve bone mass (US Patent No US 2019/0351055 A1). RJK and CWW have a patent pending on colchicine analogs to treat musculoskeletal disorders (PCT/US2018/038300). No other authors have competing interests.

## Data and Materials Availability

All data is available in the manuscript or the supplementary materials. Materials and reagents will be made available upon request, pending a signed Materials Transfer Agreement between the recipient and the University of Maryland, Baltimore.

---

**Supplemental Fig. 1.**
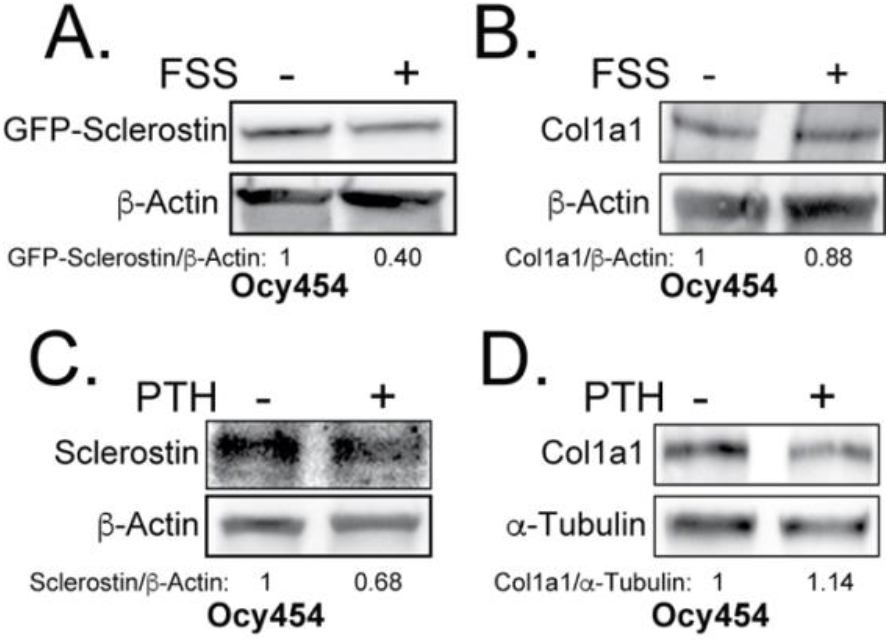
Rapid loss of sclerostin protein is specific. **(A)** Ocy454 cells transfected with GFP-sclerostin were subjected to five minutes of FSS at 4 dynes/cm^2^ and lysed immediately post-flow. Western blots were probed for sclerostin and β-Actin. **(B)** Ocy454 cells were subjected to five minutes of FSS at 4 dynes/cm^2^ and lysed immediately post-flow. Western blots were probed for pro-collagen type 1α1 and β-Actin. **(C, D)** Ocy454 cells were treated with PTH (10nM) for 30 minutes and lysed. Western blots were probed for sclerostin, β-Actin, pro-collagen type 1α1, and α-Tubulin.

**Supplemental Fig. 2.**
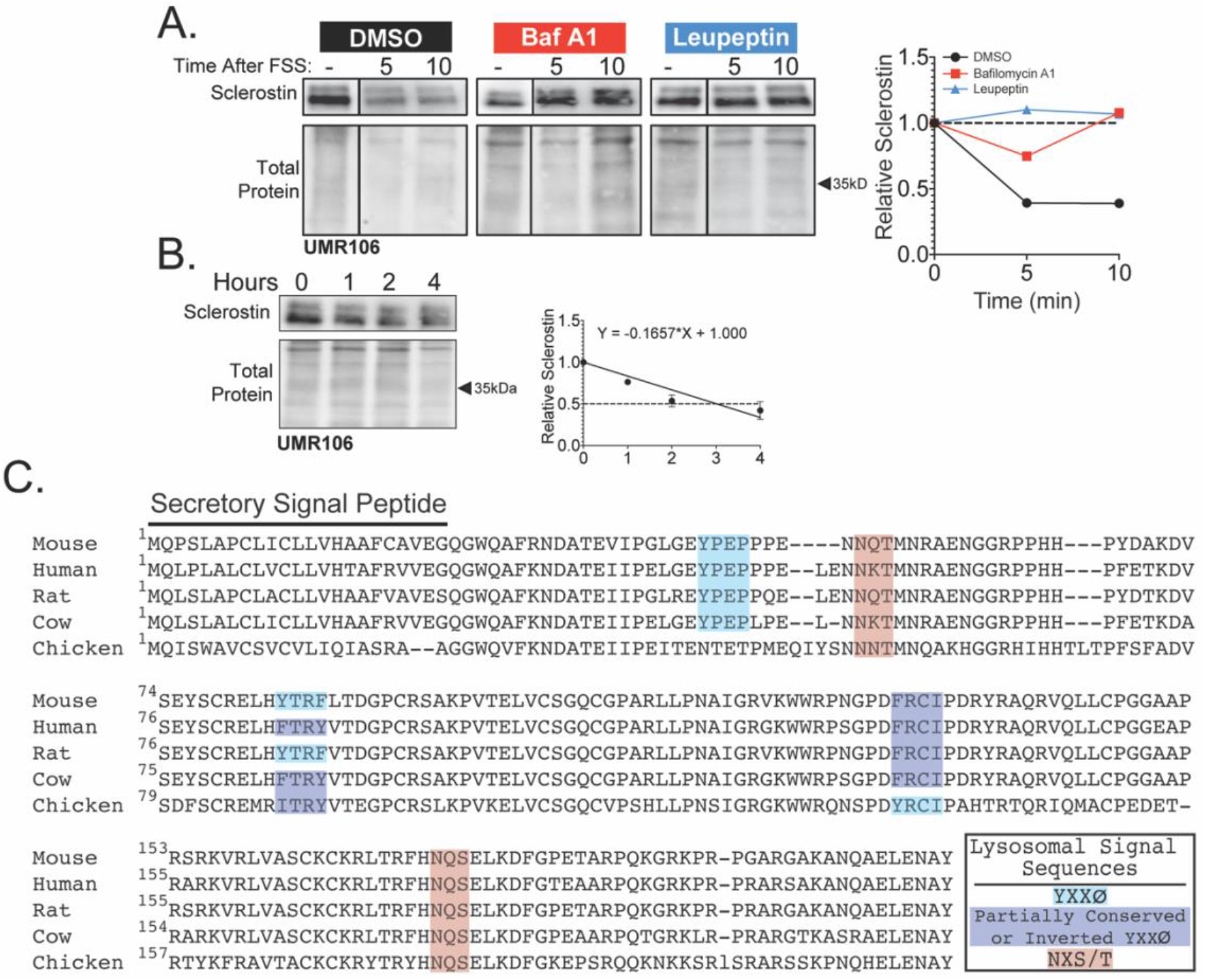
Sclerostin is rapidly degraded by the lysosome. **(A)** UMR106 cells were treated with cycloheximide (150µg/mL) and either DMSO (0.1%), Bafilomycin A1 (100nM) for 6 hours, or Leupeptin (200μM) for 4 hours prior to FSS. Cells were subjected to 1 minute of FSS at 4 dynes/cm^2^ and were lysed 5 or 10 minutes after the conclusion of FSS. Western blots were probed for sclerostin. Sclerostin abundance relative to total protein was quantified. For each antibody, blots are from a single gel and exposure; a vertical black line indicates removal of irrelevant lanes. **(B)** UMR106 cells were treated with cycloheximide (150µg/mL) and lysates were collected at 0, 1, 2, and 4 hours after treatment in the absence of stimuli. Whole cell lysates were western blotted for sclerostin abundance (n=2). Sclerostin abundance relative to the total protein was quantified. Graph shows mean ± SD and best fit linear regression constrained through y=1. **(C)** Full amino acid sequences for sclerostin from mouse, human, rat, cow, and chicken were aligned using NCBI COBALT. Putative lysosomal signal sequences are annotated. The secretory signal peptide is annotated for amino acids 1-23.

**Supplemental Fig. 3:**
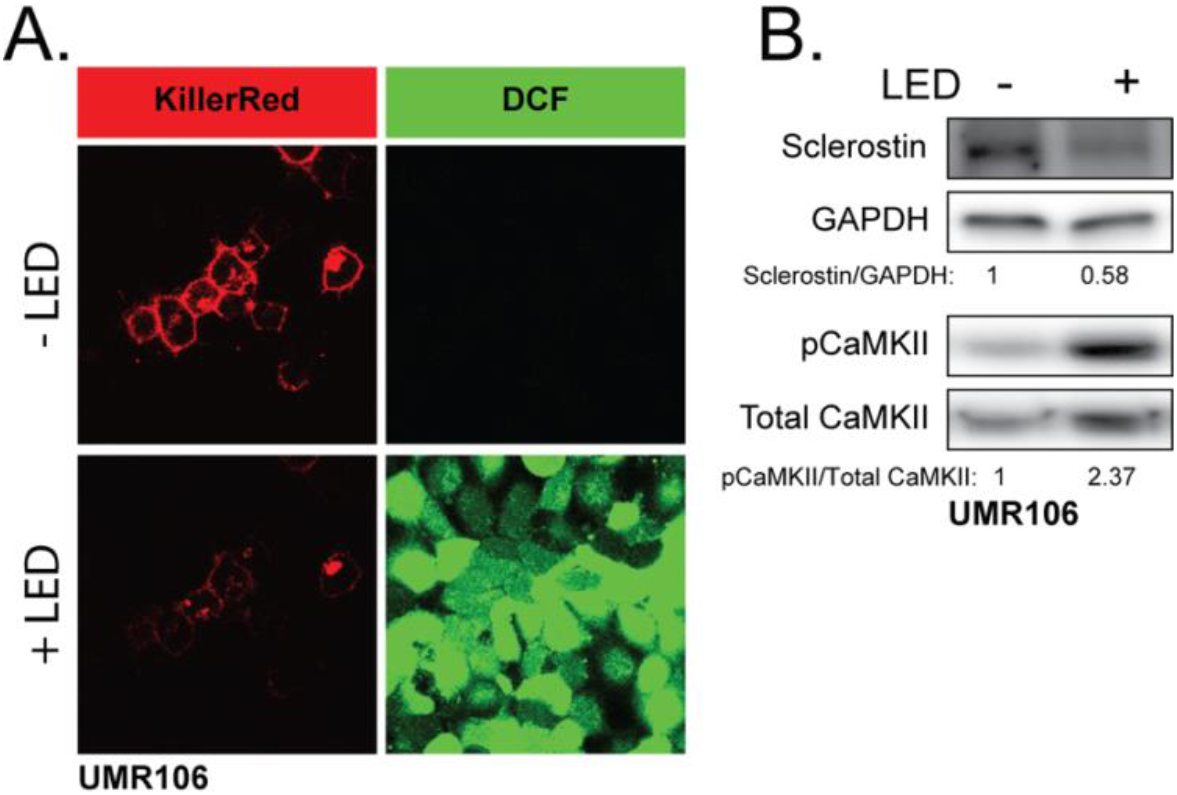
ROS is sufficient to drive CaMKII activation and loss of sclerostin protein. **(A)** UMR106 cells transfected with KillerRed imaged before and after stimulation with LED light. DCF was used to track ROS production. **(B)** UMR106 cells transfected with KillerRed were stimulated with LED light for 5 minutes and lysed 5 minutes after. Westerns were probed for sclerostin, GAPDH, pCaMKII, and total CaMKII.

